# Unravelling effects of fine-scale changes within wild-bird flocks on sociality

**DOI:** 10.1101/2023.12.05.569701

**Authors:** C.A. Troisi, J.A. Firth, S.J. Crofts, G.L. Davidson, M.S. Reichert, J.L. Quinn

**Author notes:** **Conflict of Interest:** The authors declare no conflict of interests. **Funding** This research was funded by the European Research Council under the European Union’s Horizon 2020 Programme (FP7/2007-2013)/ERC Consolidator Grant ‘EVOLECOCOG’ Project No. 617509, awarded to J.L.Q., and by a Science Foundation Ireland ERC Support Grant 14/ERC/B3118 to J.L.Q. C.A.T is funded by a Marie Sklodowska-Curie Action fellowship ‘UrbanCog’ Project No. 101062662 under the European Union’s Horizon Europe Programme. J.A.F was supported by grants from NERC (NE/S010335/1 & NE/V013483/1) and BBSRC (BB/S009752/1). **Authors contribution** Conceptualization: J.L.Q., M.S.R., J.A.F., G.L.D and C.A.T.; Methodology: J.L.Q., J.A.F, M.S.R., G.L.D., and C.A.T.; Field Work: J.A.F, S.J.C. and M.S.R.; Data curation: C.A.T and J.A.F.; Formal analysis: C.A.T. and J.A.F.; Visualization: C.A.T and J.L.Q. Writing – original draft: C.A.T with J.L.Q.; Writing – review and editing: C.A.T., J.A.F, S.J.C, G.L.D, M.S.R and J.L.Q.; Supervision: J.L.Q.; Funding acquisition: J.L.Q.; Project administration: J.L.Q.; All authors gave final approval for publication. **Data availability** Data and code are available on OSF: https://osf.io/vygw3/.

## Abstract

1. Social structure and individual sociality impact a wide variety of behavioural and ecological processes. Although it is well known that changes in the physical and social environment shape sociality, how perturbations govern sociality at a fine spatial scale remains poorly understood. By applying automated experimental treatments to RFID-tracked wild great tits (*Parus major*) in a field experiment, we examined how individual social network metrics changed when food resources and social stability were experimentally manipulated at the within-group spatial scale.
2. First, we examined how individual sociality responds when food resources changed from a dispersed distribution (50m apart) to a clustered distribution (1m apart). Second, we tested how sociality changed when individuals were restricted to feeding in a manner that mimics assortative behaviour within flocks. Third, we tested the effects of experimentally manipulating the stability of these social groupings. Finally, we returned the feeders to the original dispersed distribution to test whether effects carried over.
3. Repeatability analyses showed consistent differences among individuals in their social phenotypes across the various manipulations; dyadic association preferences also showed consistency. Nevertheless, average flock size and social centrality measures increased after the food was clustered. Some of these metrics changed further when birds were then forced to feed from only one of the five clustered feeders. There was some support for group stability at individual feeders also impacting individual social network metrics: increase in flock size was more pronounced in the stable than the unstable group. Most of the differences in sociality were maintained when the food distribution returned to the dispersed pattern, and this was caused primarily by the change in resource distribution rather than the social manipulation.
4. Our results show that perturbations in the access to resources and social group stability can change sociality at a surprisingly fine spatial scale. These small-scale changes could arise through a variety of mechanisms, including assortative positioning within groups due to, for instance, similarity among individuals in their preferences for different resource patches. Our results suggest that small-scale effects could lead to social processes at larger scales and yet are typically overlooked in social groups.

## Introduction

Social interactions have important and diverse consequences for individuals and for populations. These include effects on disease transmission, mating partner choice, access to shared information, the spread of innovations, and patterns of selection among many others (Cantor et al., 2021; Cheney et al., 2016; Ellis et al., 2019). Resource distribution is a major driver of social network structure (Beck et al., 2011; Foster et al., 2012; Heinen et al., 2022; Tavares et al., 2017). For instance, more clustered food resources can increase recurring aggregation and may be linked to stronger social bonds between individuals (Tavares et al., 2017). Increasingly social network analyses are being used to understand social interactions, often revealing important effects that would otherwise go undetected in studies of individual behaviour (e.g. Godfrey et al., 2009). Social networks and individual social connections are often stable over time (Farine & Sheldon, 2019; Fisher et al., 2016; Shizuka et al., 2014; Stanley et al., 2018) and contexts (Firth & Sheldon, 2015, 2016; Lehmann & Ross, 2011). Inevitably, individual sociality and social networks are also highly plastic (e.g. Heinen et al., 2022; Proops et al., 2021), especially in fission-fusion systems. Most evidence for the stability or plasticity of social interactions comes from observational studies or from large scale manipulations (e.g. over kilometres, between groups) (but see Heinen et al., 2022). However, individual social interactions can take place at fine spatial scales within the broader social group (Wolf et al., 2007), but less is known about how these scale up to affect broader patterns of social interaction. Likewise, broad effects on social group structure can feed back on individual social interactions (Firth et al., 2016). Here we conduct experimental manipulations, in a natural population, of i) resource distribution at a fine spatial scale (within a group) and ii) social group stability, and examine their effects on individual sociality.

Food patch distribution is a large cause of variation in social interactions (Beck et al., 2011; Foster et al., 2012; Heinen et al., 2022; Tavares et al., 2017). Dispersed food patches can increase the opportunity to interact with individuals from other groups (Tavares et al., 2017), but may also require individuals to invest more time in finding food, reducing the opportunity for social interactions (Foster et al., 2012). Resource distribution is highly variable over time, which affects individual sociality and social network stability (Cantor et al., 2021; He et al., 2019). All of these effects are likely scale-dependent (Levin, 1992; Wiens, 1989), and although they have been investigated from centimetres in captivity (Tanner & Jackson, 2011) to hundreds of kilometres in the wild (Beck et al., 2011; Cortés-Avizanda et al., 2011; Foster et al., 2012; Tavares et al., 2017), generally little is known about how small-scale variation in resource distribution affects individual sociality dynamics under natural conditions.

Group membership is clearly the main driver of sociality. It follows that changes in group membership are likely to lead to changes in individual social network metrics and social structure (Shizuka & Johnson, 2020). These changes can have long-lasting effects on individual social metrics - for instance, in macaques, the absence of policing after the loss of key male individuals led the remaining members of the group to have smaller, less diverse and less integrated networks (Flack et al., 2006) - and can also impact functional behaviour (Carter & Wilkinson, 2015; Ebensperger et al., 2016, 2017; Gazda et al., 2005; Maldonado-Chaparro et al., 2018). At the same time, individual social network positions can remain remarkably stable across years even with population turnover (Aplin et al., 2015; Farine & Sheldon, 2019; Shizuka et al., 2014) and when individuals lose close associates (Boucherie et al., 2017; Firth et al., 2017).

In all of these group membership studies, however, changes in individual sociality in the group are perhaps inevitable, since typically individuals were removed or added to social groups in the experimental manipulations (Boucherie et al., 2017; Firth et al., 2017; Maldonado-Chaparro et al., 2018). Behavioural changes within groups of constant membership could also lead to changes in who individuals interact with. For example, an individual that develops a new innovative behaviour (Kulahci & Quinn, 2019; Wascher et al., 2018) can become more central in the group (Kulahci et al., 2018). The development of persistent assortative interactions among individuals within groups - due to, for instance, similarity among individuals in their preferences for different resource patches (Caillaud & Via, 2000; Crook, 1999; Martin, 2013; Snowberg & Bolnick, 2008), or in their preferred positions within groups linked to predation risk (Heathcote et al., 2017; Lambert et al., 2021) - could similarly feed back onto individual social network metrics and social structure generally. However, to date this latter possibility has not been tested experimentally.

In this study we manipulated fine-scale resource distribution and social stability in great tits (*Parus major*). Great tits form fission-fusion flocks during the non-breeding season in woodland habitat and readily come to feeders where their behaviour can be automatically detected using passive integrated transponders (Aplin et al., 2013; Cauchoix et al., 2022; Cooke, 2021; Reichert et al., 2020). We estimated flock sizes and individual social centrality - among the most important descriptors of network positioning. First, we describe patterns of visitation over time and use repeatability analyses (Stoffel et al., 2017) to test whether our social centrality measures captured consistent behaviour, i.e., intrinsic differences among individuals in their sociability. We then explore our four main hypotheses. First, we tested whether fine-scale variation in resource distribution affects individual social network metrics (SNM). We expected that by moving feeders closer together from an initially dispersed to a clustered treatment, flock sizes and individual social connectedness should increase. Second, we tested whether forcing groups of individuals to use specific feeders in the clustered 5-feeder array, mimicking, for example, assortative patch use or assortative positioning within flocks, modified individual sociality. We predicted that forcing individuals to forage at a specific feeder might disrupt the connections previously formed, and as such their flock sizes and individual social connectedness would decrease. Third, we tested whether individual social network metrics were affected by social stability, where groups of individuals were allocated to one of the five feeders for two additional phases, either with the same individuals in each group across each phase (stable treatment), or with a random selection of new individuals (unstable). We predicted that, at the end of the treatment, individuals in the stable treatment would have stronger ties with fewer individuals, because they had more opportunities to interact repeatedly with the same individuals, compared to those who shared feeders with different individuals across each of the three phases and therefore more individuals over the three phases cumulatively. Finally, we tested whether the effects of our manipulations collectively were transient or if they would persist in a different context by returning the feeders to the original dispersed distribution. On the one hand, because great tits live in a fission-fusion society and are adapted to regular small-scale changes in their (physical and social) environment, any effects observed during the manipulations might be expected to be transient, in which case we expected that individuals would associate with the same individuals at the start and the end of the experiment, and individual social network metrics would revert back to their initial value. On the other hand, if prolonged associations between individuals have longer term carry-over effects, we predicted that the manipulations observed would be persistent, and especially so in the stable treatment, where individuals had more opportunity to interact with the same individuals and create stronger bonds.

## Methods

### Study site and species

The study took place in Wytham Woods, Oxford, UK. Great tits and blue tits (*Cyanistes caeruleus*) were fitted with PIT tags following Reichert et al. (2020). During the winter, these birds form fission-fusion foraging flocks and move around the woodland (Farine et al., 2015; Firth & Sheldon, 2016).

Data collection took place during the winter season from November 2017 to February 2018. Four sites were used early in the season (November-December 2017), and four different sites were used later in the season (January-February 2018). Only data from great tits were used in this study because a very high proportion of the population is tagged: an estimated ∼80-90% of great tits were tagged at this time based on previous studies with similar trapping effort (Aplin et al., 2013; Matechou et al., 2015) and 258 individuals used our feeders (see Table S1 for age and sex profiles following the STRANGE recommendations (Webster & Rutz, 2020)).

### Treatments

Feeders containing sunflower seeds and equipped with an RFID antenna were active each day during daylight hours from 0700 to 1630, and PIT tagged birds had ad libitum access to food, at some or all of these feeders, during those times (see Reichert et al., 2020 for details). At each of the eight sites, feeders were arranged according to four main treatments: the initial dispersed, the open clustered, the assortative clustered, and the final dispersed treatment (see Figure 1.A). Note that we use the word “assortative” in the sense that restricting individual access to individual feeders in effect placed them into groups of individuals that shared a feeder, even if they had not been grouped based on any a priori shared characteristic. The initial and final dispersed treatments consisted of one feeder at each of two locations that were 50m apart, during which all birds could feed from either feeder. The two clustered treatments were sandwiched in time between the two dispersed treatments, and consisted of five feeders 1m apart at one location. We acknowledge that the availability of food could also have been higher in the clustered treatment, and therefore the treatment may reflect a combination of resource distribution and food availability. However, we think the distribution was the dominant effect because i) the number of individuals that visited the feeders was lower in the clustered treatment than in the previous initial dispersed treatment (see below); ii) food availability was likely not a limiting factor because seeds were always available from all allocated feeders and delivery of the single seed reward was effectively instantaneous, notwithstanding any queuing that took place around the feeders.

**Figure 1:**
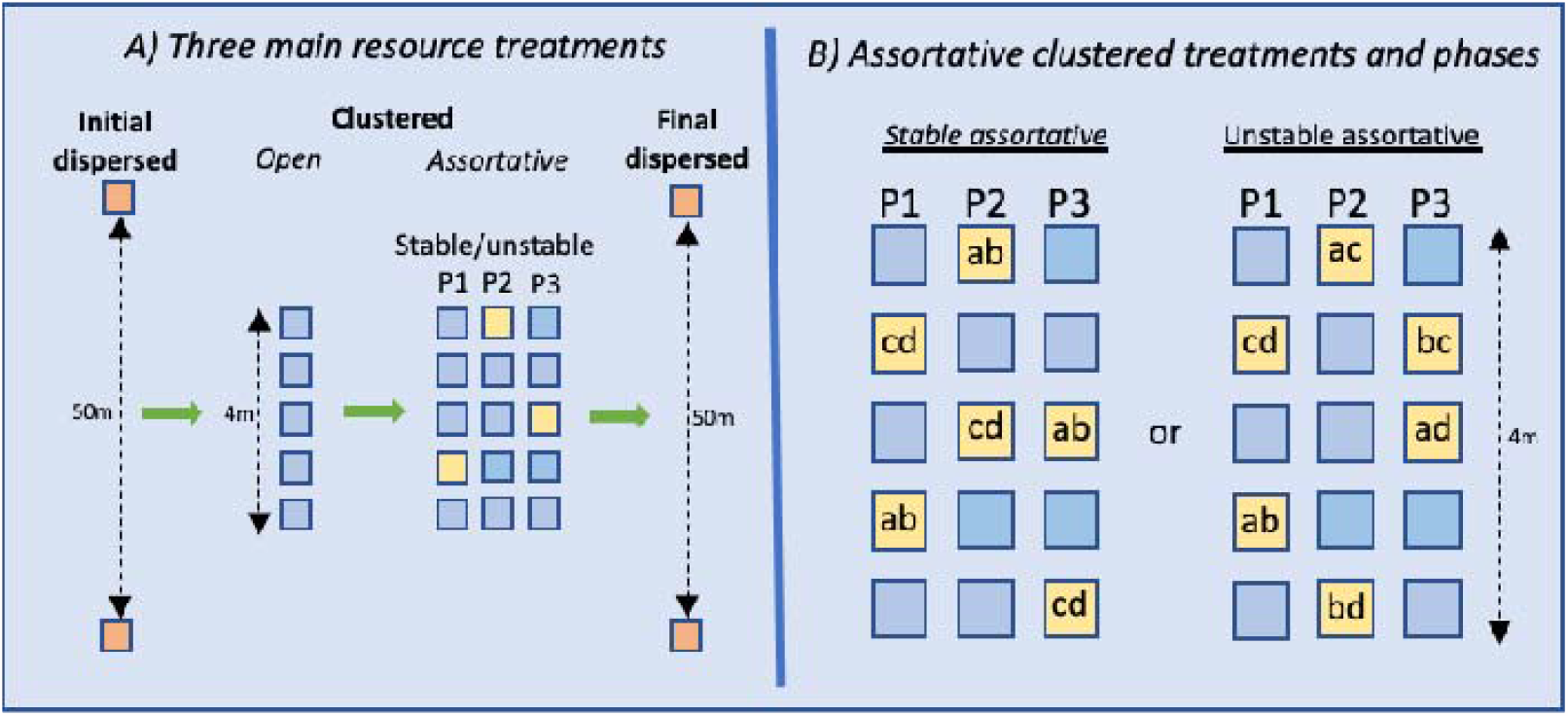
A) Layout of the experiment showing the food distribution across the three main treatments where food distribution was manipulated (initial dispersed; clustered; final dispersed). The clustered treatment was further split into open clustered and assortative clustered subtreatments, and B) the assortative treatment in turn was further split into stable and unstable subtreatments, each of which ran over three phases (P1; P2; P3)

In the open clustered treatment, birds could obtain food from any of the 5 feeders. In the assortative clustered treatment, birds could only access one feeder, to which they were randomly allocated. If birds landed on their assigned feeder, the feeder would open and they had access to the food. If they landed on any of the 4 other feeders, the feeder would remain closed, and the bird would not have access to the food, but its visit would still be recorded feeding into the social network metrics. Great tits quickly learned which feeder they were allocated to, usually within the first day and usually after less than 30 visits (Reichert et al., 2020).

The assortative clustered treatment further consisted of 3 phases (see Figure 1.B). After eight to ten days (eight days in 2017 and ten days owing to operational differences in 2018) of the initial feeder assignment in the assortative clustered treatment, we manipulated social stability by randomly allocating birds to a different feeder, either with the same individuals as in their original feeder assignment (the stable assortative clustered treatment; run at four sites) or with a random selection of predominantly different individuals (the unstable assortative clustered treatment; run at the remaining four sites). After another eight to ten days we then repeated the feeder reassignment procedure; birds in the stable treatment were again reassigned along with the same individuals from their original feeder assignment and birds in the unstable treatment were again reassigned with a new randomly selected group of individuals.

The raw dataset consisted of rows containing the date, time and PIT tag for each detected visit at each feeder. We considered consecutive detections of the same bird to the same feeder within 2s of each other to be a single visit (following Evans et al., 2018; and Reichert et al., 2020).

### Individual network metrics

Using the spatio-temporal data of visits to feeders, “flocks” (or ‘flocking events’) were identified at each location using a machine learning algorithm (Psorakis et al., 2012, 2015). A Gaussian mixture model assigned each individual visit from each bird to the flocking event for which it had the highest probability of belonging, without imposing assumptions about the temporal boundaries of flocks (Psorakis et al., 2012). This allowed us to calculate an average flock size for each individual at each location, during each treatment, or each phase of the clustered treatment, separately.

Edges in the social network were assigned to each individual appearing in the same flocking event. For each possible pair of individuals, we then counted the number of flocking events in which both individuals were present. We used these data to quantify the association strength for each dyad as a “simple ratio index”: the number of times both individuals were seen in the same flocking events ÷ (the number of times individual A was seen in a flocking event without B + the number of times individual B was seen in a flocking event without A + the number of times individuals A and B were both seen in the same flocking event) (Cairns & Schwager, 1987; Whitehead, 2008).

For each individual, we also calculated two commonly used social network centrality metrics: 1) weighted degree - the sum of all the focal individual’s weighted associations (i.e. the number of times each association between two individuals was observed) with all other individuals (also known as node ‘strength’); and 2) weighted eigenvector centrality - a measure of the total amount of social associations of an individual’s associates (i.e. the centrality of their flockmates); for instance, an individual that associates with highly sociable individuals would have high eigenvector centrality whilst an individual that associates with peripheral individuals would have low eigenvector centrality. As such, these network metrics represent a range of measures of individual centrality (Albery et al., 2020) on an increasing scale of complexity.

### Ethics

We performed the experiment in accordance with the Association for the Study of Animal Behaviour ethical guidelines, under permission of Oxford University Internal Animal Welfare Committee (Zoology), and the Animal Experimentation Ethics Committee of the University College Cork. The Health Products Regulatory Authority approved the ethics for the project number AE19130/P017. All bird ringing and tagging was carried out under standard licencing permissions from the British Trust for Ornithology (BTO).

### Data analysis

All analyses were conducted in R version 4.0.2 (R Core Team, 2020). The package *ggplot2* was used for plotting graphs (Wickham, 2016).

#### General analyses

Initially we tested whether our measures of individual sociality captured intrinsic differences among individuals in social behaviour by estimating the repeatability of flock sizes (for each individual, averaged across all flocking events within a treatment) and social network metrics (calculated for each treatment separately) across the experiment, resulting in 6 measures per individual (Figure 1). We included only birds that appeared in all 6 treatment levels (N=70; 68 individuals, with 2 individuals present at two sites). We used the *rptR* package to calculate repeatability (Stoffel et al., 2017), using lmms with a Gaussian error distribution.

To provide context for the main analyses, and to test whether temporal variation in feeder usage over the course of the experiment might confound the main hypotheses testing, we explored whether i) the number of individuals detected per treatment, ii) the number of visits per individual per day, and iii) the number of flocks an individual was found in varied across all 6 treatment levels. The first of these was analysed using a Poisson distribution, and the remaining two as Gaussian distributions, and the package *lmerTest* was used for linear mixed models (Kuznetsova et al., 2017). For the site level analysis (number of individuals per treatment) we included experimental treatment as a fixed factor and site as a random effect. For the individual level analyses (number of visits per individual and number of flocks) we included, sex, age (adult vs juvenile), and experimental treatment as fixed factors as well as individual identity and site as random effects.

#### Hypothesis testing

In all of the analyses below, we used linear mixed models with Gaussian error distribution from the *lmerTest* package (Kuznetsova et al., 2017), with separate models for each of the individual social network metrics as the dependent variable (flock size, weighted degree, eigenvector centrality). Social network based metrics necessarily violate the assumption of independence, so in all cases we also compared the model estimates to those calculated from null models using node-based permutations (Whitehead, 2008). We report p-values showing where the observed estimates fall within the distribution of estimates from the 1000 permutations for each model i.e. if p<0.05 then the observed estimate falls outside of the 95% range of the null expectation (Whitehead, 2008). A small number of birds (usually 1-3 in any one analysis) were present at several of our eight sites (as four sites were used early in the season, and 4 sites later in the season) and this was accounted for using individual as a random effect in all analyses. If they were present at two sites, they were only ever allocated to a feeder at one site during the assortative clustered treatments. In the analyses of dyadic association strength, sex was considered unimportant so the sample sizes indicated included birds of unknown sex.

##### Changing from dispersed to clustered food distribution influences social behaviour (H1)

Birds of known sex present in both initial dispersed and open clustered treatment levels were included (N = 121 individuals; N=3 individuals appeared at more than one site; Table S1). The model structure for each of the three social network metrics included individual and site as random effects, and fixed effects of resource treatment (initial dispersed vs open clustered), sex, age, the number of flocking events individuals took part in, and the number of individuals in the network at that site during that treatment. The latter two variables ensured that any observed treatment effects were not simply due to changes in general activity or the total numbers of individuals present over time. Finally, we estimated whether individuals associated with the same individuals in both treatments of the experiment using a linear mixed model, with the simple ratio index (a measure of association strength) during the open clustered treatment as the response variable and the simple ratio index during the initial dispersed treatment as a fixed effect, using the *lmerTest* package (Kuznetsova et al., 2017) (N=131 individuals; N=3 individuals appeared at more than one site). We included the identity of both individuals of each dyad and site as random effects.

##### Assortative feeding on a fine spatial scale changes social behaviour (H2)

Birds of known sex were included in these analyses only if present in both the open clustered and the first assortative clustered (P1) treatment levels (N=100 unique individuals; N=2 individuals appeared at more than one site; Table S1). Once again, three separate models were run for each social network metric and included the same random and fixed effects as for H1. We then tested whether dyadic associations during the open clustered phase predicted the assortative clustered P1 phase in the same manner as described in H1, and the sample size was N=107 individuals (2 individuals appeared at more than one site).

##### Social stability influences the effect of the assortative feeding on social behaviour (H3)

Birds of known sex were included in these analyses only if present in each of the three assortative clustered treatment phases, but the main factor used for hypothesis testing included only the first and third assortative clustered treatment phases to examine the overall effect of the assortative clustered treatment (N=91 unique individuals; N=3 individuals appeared at more than one site; Table S1). Here we tested the hypothesis using an interaction between resource treatment (assortative clustered P1 and P3) and social stability (stable vs unstable), predicting that individuals in the stable treatment would have stronger ties (higher weighted degree and weighted eigenvector centrality) with fewer individuals (smaller flock size) after the assortative treatment compared to those who shared feeders with different individuals across each of the three phases. Random and additional fixed effects included were the same as for H1, with the addition of the social stability treatment (stable vs unstable) as main effect, and the interaction between social stability treatment and resources treatment. We ran posthoc tests with the *emmeans* package to examine the effects of the resource treatment on social network metrics for birds in each social treatment (Lenth, 2019). We also examined whether the social stability treatment influenced the simple ratio index of association using a similar model to H2, but again adding the interaction between the simple ratio index during the assortative clustered P1 treatment and social stability treatment. The sample size was N= 94 individuals (3 individuals appeared at more than one site).

##### Observed changes in sociality persisted when dispersed food treatment was restored (H4)

To examine the persistence of the observed changes we compared the social metrics from the initial dispersed to the final dispersed treatment levels, controlling for sex, age, number of flocking events an individual took part in, and the number of individuals in the social network. We tested the effect of resource treatment (initial and final dispersed) as well as the interaction between resource treatment and social stability treatment (stable vs unstable) to examine whether any observed differences between stable and unstable groups at the end of the assortative clustered treatment were still observed at the final dispersed stages. In this analysis, we included only individuals of known sex that were present at all 6 stages of the experiment (N=67 unique individuals; N=2 individuals appeared at more than one site; Table S1). Posthoc tests of the interaction were carried out with the *emmeans* package (Lenth, 2019). We also tested whether the dyadic association score in the final dispersed treatment was predicted by the dyadic association score in the initial dispersed treatment, and whether the association depended on the social stability treatment by including the resource treatment, social stability treatment (stable vs unstable), and their interaction as fixed effects. We included the identity of both individuals of each dyad and site as random effects. The sample size for this analysis was N= 68 individuals (2 individuals appeared at more than one site).

## Results

All individual social metrics showed moderate repeatability, and our measures therefore capture intrinsic among-individual differences in their sociality (flock size: R = 0.353, CI = 0.232 - 0.459, p < 0.001; weighted degree, R= 0.256, CI = 0.150 - 0.359; p < 0.001; weighted eigenvector centrality, R = 0.104, CI = 0.023 - 0.190, p = 0.002). The number of individuals at each site was significantly lower in the open clustered and final dispersed treatment than in the initial dispersed treatment, but remained similar between the initial dispersed and the three assortative clustered treatments (Table S2; Figure S1a). The number of visits each bird made per day remained similar between the initial dispersed and open clustered treatments, but birds made significantly fewer visits per day in the initial dispersed than in the three assortative clustered and the final dispersed phases (Table S2; Figure S1b). The number of flocking events per individual per day remained similar between the initial dispersed and open clustered treatments, but individuals took part in significantly fewer flocks per day in the initial dispersed than in the three assortative clustered and the final dispersed phases (Table S2; Figure S1c). Thus, for each treatment level in all further analyses, we controlled for the number of individuals at sites and the number of flocks it took part in. We did not include the number of visits each bird made because it was strongly colinear with the number of flocking events it took part in (r=0.911).

### From dispersed to clustered (H1)

All measures of sociality were significantly higher in the open clustered treatment than in the preceding initial dispersed treatment (Figure S2; Table S3; null model tests Table S4). Dyadic associations in the initial dispersed treatment predicted associations in the open clustered treatment (B = 0.43 (95%CI =0.358-0.497); intercept B = 0.073 (95%CI = 0.046-0.099); N = 1271 pairs and 131 unique individuals; Figure 2a).

**Figure 2:**
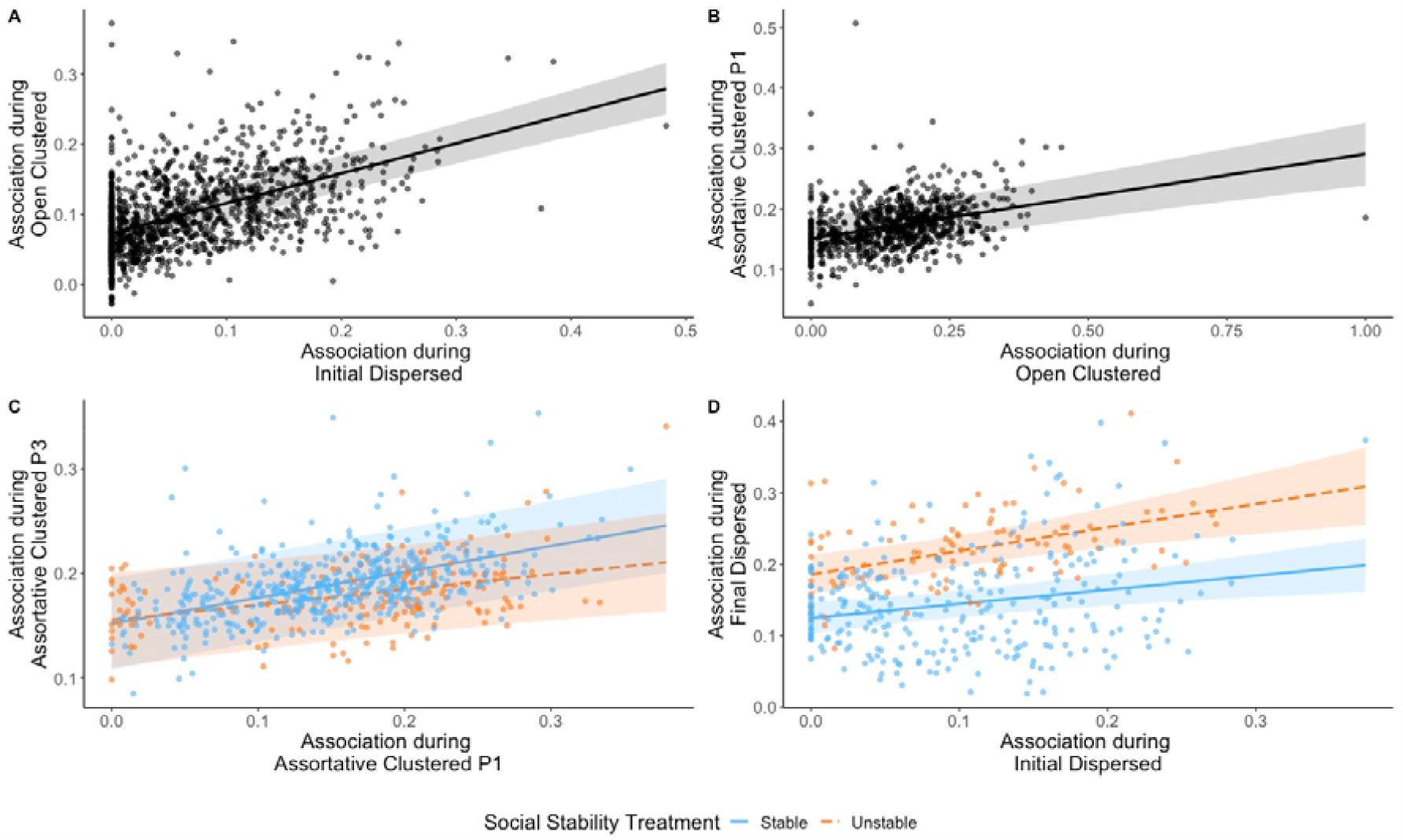
Partial residual plots showing how pairwise associations in one treatment predicted these in the next for: a) the initial dispersed treatment and the open clustered treatment; b) the open clustered treatment and the assortative clustered treatment (P1); c) the assortative clustered P1 treatment and P3 treatment; and d) the initial dispersed and final dispersed treatments. For c) and d), separate lines are shown for birds in the stable (blue) and unstable (orange) social stability treatments. Interaction was ns. for c) and d). Shaded areas are the 95% confidence intervals from corresponding models in the main text and in Table S7 and S10. We added random effects for the identity of both individuals of each dyad and site.

### From open clustered to assortative clustered (H2)

Restricting individuals to being able to access food from only one of the five feeders in the array led to a significant increase in flock size, and a significant decrease in weighted degree and weighted eigenvector centrality (open clustered vs. assortative clustered P1 treatment level; Table 1, Figure 3). Comparison to the null models gave qualitatively similar results (Table S5).

**Table 1:** Linear mixed models of how each of four individual social network metrics changed from the open clustered to the assortative clustered treatments. Site and individual identity were included as random effects. ^1^ baseline = female; ^2^ baseline = adult; ^3^ baseline = open clustered

| Dependent variable | Independent variables | Estimate (SE) | 95% CI | P value |
| --- | --- | --- | --- | --- |
| Flock Size | Intercept | 1.35 (0.504) | 0.391; 2.30 | 0.011 |
|  | Flocking events | 0.001 (0.002) | -0.002; 0.004 | 0.565 |
|  | Individuals in local network | 0.180 (0.018) | 0.145; 0.216 | <0.001 |
|  | Sex (male)<sup>1</sup> | -0.089 (0.115) | -0.313; 0.135 | 0.441 |
|  | Age (juveniles)<sup>2</sup> | -0.030 (0.125) | -0.273; 0.212 | 0.808 |
|  | Resource (assortative clustered P1)<sup>3</sup> | 0.765 (0.217) | 0.346; 1.18 | <0.001 |
| Weighted Degree | Intercept | -0.318 (0.310) | -0.917; 0.279 | 0.311 |
|  | Flocking events | 0.018 (0.001) | 0.016; 0.020 | <0.001 |
|  | Individuals in local network | 0.085 (0.012) | 0.063; 0.107 | <0.001 |
|  | Sex (male)<sup>1</sup> | 0.010 (0.071) | -0.129; 0.148 | 0.891 |
|  | Age (juveniles)<sup>2</sup> | -0.069 (0.077) | -0.219; 0.081 | 0.374 |
|  | Resource (assortative clustered P1)<sup>3</sup> | -1.02 (0.134) | -1.28 -0.762 | <0.001 |
| WEVC | Intercept | 0.567 (0.071) | 0.433; 0.702 | <0.001 |
|  | Flocking events | 0.005 (0.0003) | 0.005; 0.006 | <0.001 |
|  | Individuals in local network | -0.004 (0.003) | -0.009; 0.002 | 0.190 |
|  | Sex (male)<sup>1</sup> | 0.001 (0.022) | -0.043; 0.043 | 0.955 |
|  | Age (juveniles)<sup>2</sup> | -0.053 (0.024) | -0.100; -0.006 | 0.031 |
|  | Resource (assortative clustered P1)<sup>3</sup> | -0.371 (0.039) | -0.445; -0.295 | <0.001 |

**Figure 3:**
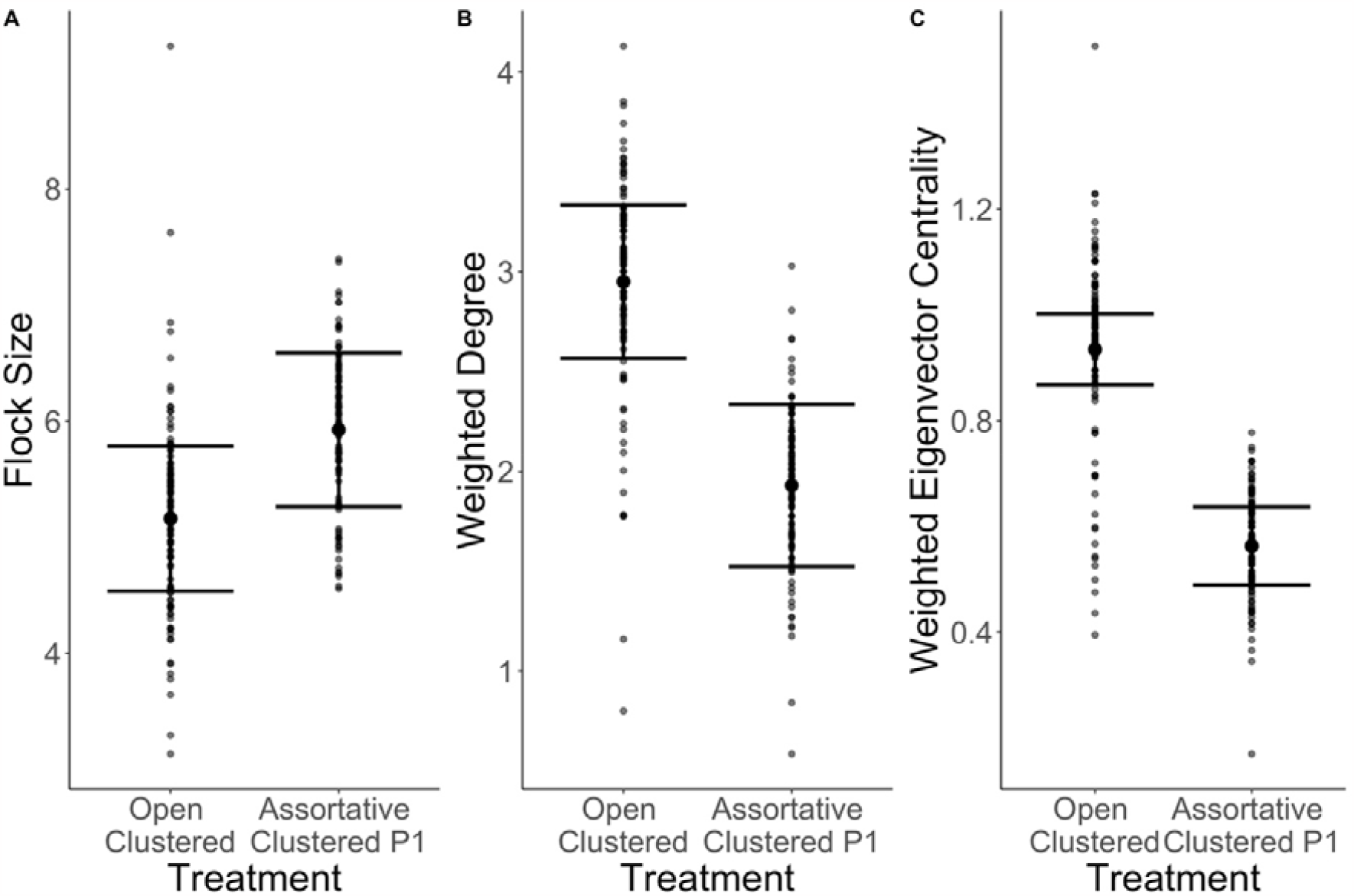
Partial residual plots showing changes in (A) flock size, (B) weighted degree and (C) weighted eigenvector centrality, across the open clustered and assortative clustered P1 treatment. Error bars are 95% confidence intervals based on the linear mixed models in Table 1.

Once again, social preferences during the open clustered treatment predicted those in the assortative clustered P1 treatment (B = 0.139 (95% CI = 0.092-0.187); Intercept: B = 0.151 (95% CI = 0.117-0.188); N pairs = 800; N unique individuals = 107; Figure 2b).

### Social stability (H3)

Social stability from the first to the third assortative clustered phases (P1-P3) significantly influenced the effect of resource treatment on flock size (Resource × social stability effect; Table 2; Table S6, Figure 4), and again the null models gave qualitatively similar results (Table S7). A significant increase in flock size was more pronounced in the stable group than in the unstable group (Table S6; Figure 4a). Weighted degree significantly increased across the assortative clustered phases, but this increase in weighted degree was similar for both social stability treatments (Table S6, Figure 4b). There was weak (non-significant) support for the stability treatment influencing weighted eigenvector centrality, which decreased more for the stable than for the unstable treatment (Table S6, Figure 4c)

**Table 2:** Linear mixed models of how changes in each of four individual social network metrics, from the start (P1) to the end (P3) of the assortative clustered resource treatment level, were influenced by social stability, as tested by their interaction. Site and individual identity were included as random effects. ^1^ baseline = female; ^2^ baseline = adult; ^3^ baseline = assortative clustered P1; ^4^ baseline = stable

| Dependent variable | Independent variables | Estimate (SE) | 95% CI | P value |
| --- | --- | --- | --- | --- |
| Flock Size | Intercept | 0.946 (0.507) | 0.006; 1.88 | 0.082 |
|  | Flocking events | -0.0005 (0.001) | -0.002; 0.001 | 0.530 |
|  | Individuals in local network | 0.255 (0.021) | 0.216; 0.293 | <0.001 |
|  | Sex (male)<sup>1</sup> | -0.017 (0.075) | -0.158; 0.125 | 0.811 |
|  | Age (juveniles)<sup>2</sup> | 0.040 (0.089) | -0.158; 0.125 | 0.626 |
|  | Resource (assortative clustered P3)<sup>3</sup> | 1.02 (0.097) | 0.830; 1.21 | <0.001 |
|  | Social stability (unstable)<sup>4</sup> | 0.199 (0.548) | -0.844; 1.25 | 0.731 |
| | Resource (assortative clustered P3) × social stability (unstable) | -0.596 (0.153) | -0.885; -0.305 | <0.001 |
| Weighted Degree | Intercept | 0.124 (0.302) | -0.465; 0.696 | 0.690 |
|  | Flocking events | 0.012 (0.0004) | 0.011; 0.013 | <0.001 |
|  | Individuals in local network | 0.042 (0.012) | 0.019; 0.069 | 0.001 |
|  | Sex (male)<sup>1</sup> | 0.039 (0.0423) | -0.043; 0.122 | 0.361 |
|  | Age (juveniles)<sup>2</sup> | 0.044 (0.047) | -0.048; 0.135 | 0.354 |
|  | Resource (assortative clustered P3)<sup>3</sup> | 0.232 (0.057) | 0.120; 0.341 | <0.001 |
|  | Social stability treatment (unstable)<sup>4</sup> | -0.350 (0.334) | -0.976; 0.260 | 0.346 |
| | Resource (assortative clustered P3) × social stability (unstable) | -0.007 (0.087) | -0.173; 0.165 | 0.934 |
| WEVC | Intercept | 1.09 (0.123) | 0.859; 1.33 | <0.001 |
|  | Flocking events | 0.004 (0.0001) | 0.003; 0.004 | <0.001 |
|  | Individuals in local network | -0.042 (0.004) | -0.050; -0.032 | <0.001 |
|  | Sex (male)<sup>1</sup> | 0.063 (0.015) | -0.022; 0.035 | 0.672 |
|  | Age (juveniles)<sup>2</sup> | -0.007 (0.016) | -0.039; 0.025 | 0.683 |
|  | Resource (assortative clustered P3)<sup>3</sup> | -0.099 (0.020) | -0.138; -0.061 | <0.001 |
|  | Social stability (unstable)<sup>4</sup> | -0.184 (0.145) | -0.460; 0.087 | 0.262 |
| | Resource (assortative clustered P3) × social stability (unstable) | 0.049 (0.030) | -0.009; 0.108 | 0.108 |

**Figure 4:**
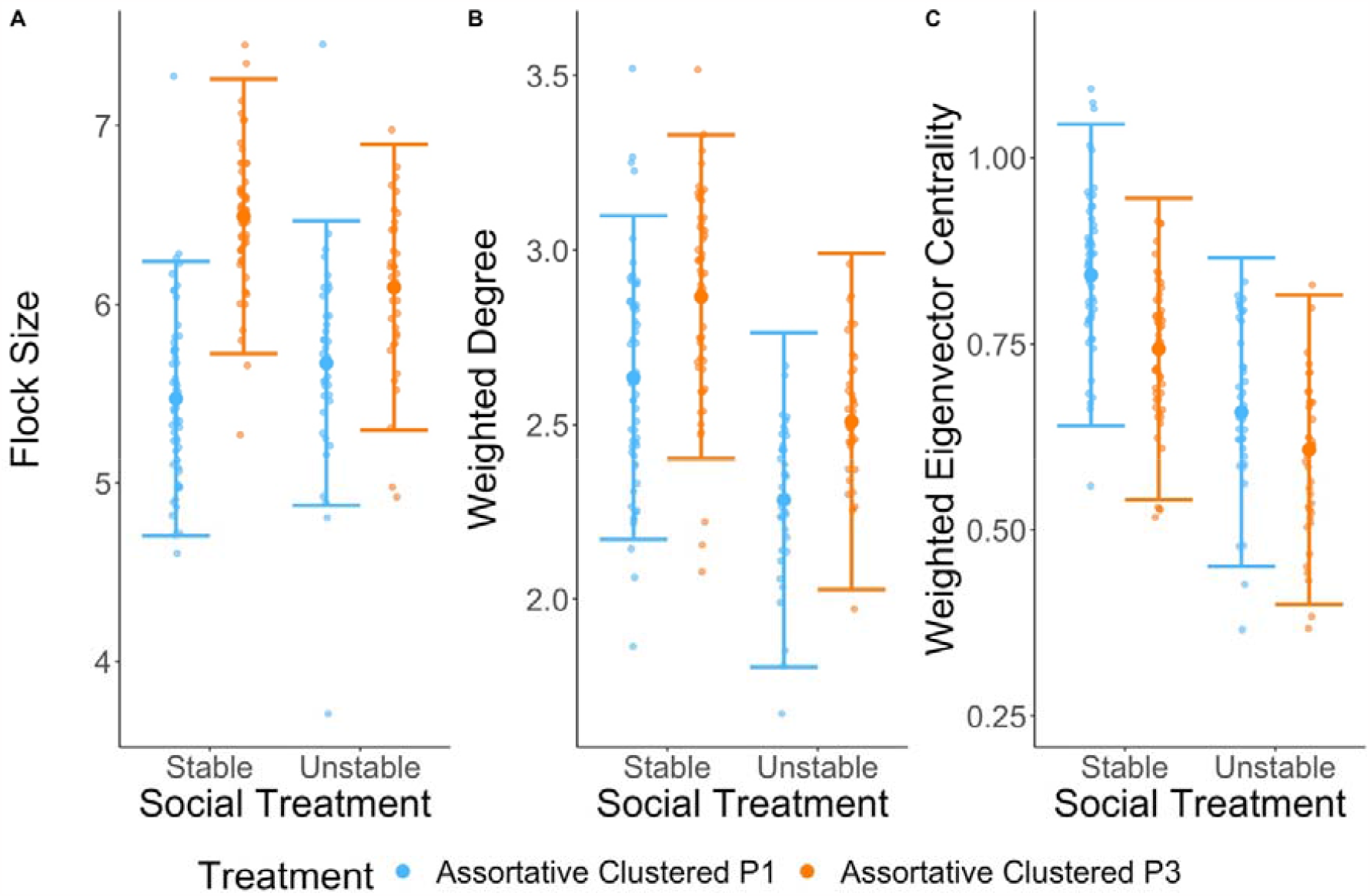
Plots showing the partial residuals for (A) flock size, (B) weighted degree and (C) weighted eigenvector centrality, and how these changed across the assortative clustered P1 and assortative clustered P3 treatment for each social stability treatment. Error bars are 95% confidence intervals based on models in Table 3.

Dyadic associations during the assortative clustered P1 treatment predicted the associations during the assortative clustered P3 treatment (Table S8; Figure 2c; N pairs = 670; N unique individuals = 94). There was weak (non-significant) support for this correlation being stronger in the stable than in the unstable treatment (Association during P1 × social stability, B ± SE = -0.101 ± 0.065, P = 0.121; Figure 2c).

### Persistence of effects: initial vs final dispersed treatment level (H4)

All three social network metrics were higher in the final dispersed treatment level than they had been in the initial dispersed treatment level, even after controlling for changes in network membership and increases in visit rates (main effects of treatment in Table S9 for mixed models, and Table S10 for null models; Figure 5). The increase in flock size was significantly greater for the unstable than the stable treatment level; there was no evidence that stability significantly affected the change in weighted degree or weighted eigenvector centrality from initial to final dispersed stages (Resource × stability treatment effects in Table S9; Table S11; Figure 5).

**Figure 5:**
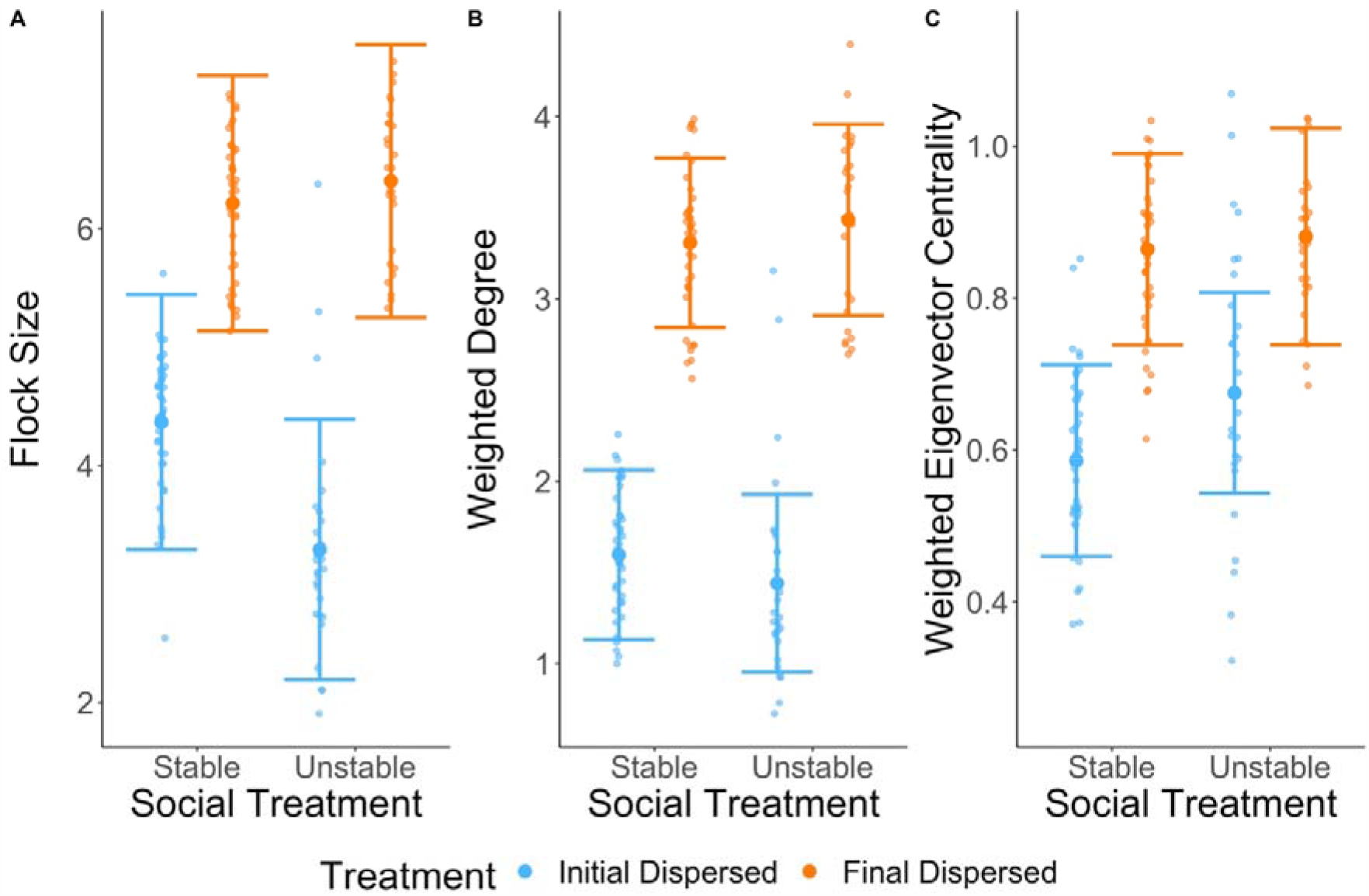
Plots showing the partial residuals for (A) flock size, (B) weighted degree and (C) weighted eigenvector centrality, across the initial dispersed and final dispersed treatment, for each social stability treatment. Error bars are 95% confidence intervals based on models in Table S8.

Dyadic associations during the initial dispersed treatment level predicted those in the final dispersed treatment level (B=0.228 (95% CI: 0.125; 0.335); Intercept B=0.157 (95% CI: 0.120-0.196); N pairs = 379; N unique individuals = 68; Figure 2d). There was no evidence that the social stability treatment significantly influenced the relationship between dyadic associations at the two time points (association initial dispersed× social stability, B ± SE = 0.132 ± 0.109, P = 0.229; Table S12, Figure 2d). During the final dispersed treatment level, dyadic associations were stronger for birds that experienced the unstable treatment than for birds that experienced the stable treatment (B=0.074 (95% CI: 0.037-0.110); Intercept B=0.143 (95% CI: 0.118-0.169). This was not the case during the initial dispersed treatment level (B=0.002 (95% CI: -0.054-0.057); Intercept B=0.097 (95% CI: 0.059-0.134)).

## Discussion

Individuals had repeatable social network metrics, and were consistent in whom they associated with throughout the experiment. However, we found that at a small scale, changes in both food distribution and social stability influenced individual level social network metrics. Some of these effects lasted even when food distribution and social groupings were reverted back to their original structure.

### Resource distribution and individual sociality (H1 & H2)

Manipulating resource distribution from two feeders 50m apart (dispersed) to a clustered array of five feeders only 1m apart led to an immediate increase in individual sociality for all metrics, in line with previous studies (Tanner & Jackson, 2011; Tavares et al., 2017; Zahavi, 1971). This is unsurprising simply because the total numbers of birds at the site only declined by about 20% and yet suddenly the remaining birds were feeding on additional food resources at one location instead of two. The response observed, therefore, reflects the fact that when birds have to find new foraging patches they ultimately converge on similar feeding locations, likely through a variety of mechanisms linked to shared information (Ward & Webster, 2016).

Our more novel finding was that sociality changed considerably when individuals were then restricted to separate single group locations on a fine-spatial scale, even though the spatial scale and location of the food resources remained unchanged. Flock sizes increased, which was likely caused by reduced feeder access, forcing individuals to spend more time at the location to get the food (presumably being forced to queue for longer, and/or learning which feeder provides food) and being registered in the same flocks. As predicted, the same manipulation reduced both the strength of connections, as indicated by declines in weighted degree, and individuals’ overall social connectedness, as indicated by a reduction in weighted eigenvector centrality. Individuals occurred in larger but less well-connected flocks. This decrease in social connectedness could imply a strong trade-off between ensuring access to resources (without being able to rely on social information) - which may have required individuals to stay at feeders to attempt to get food and in the process overlapping with many more individuals but having less reliable connections to specific individuals - and maintaining strong connections. To our knowledge this is the first experimental demonstration that restricting where individuals feed at a spatial scale smaller than that encompassed by a single social ostensible group, can change the sociality of individuals, demonstrating the importance of scale in understanding the effects of resource use on social interactions (Cortés-Avizanda et al., 2011; Johnson et al., 2002).

Constraints on where individuals feed within group foraging locations can arise through a variety of mechanisms, for example because of competitive ability when patch quality varies, risk taking behaviour when risk varies and personality (Quinn et al., 2012; Webster & Ward, 2011). Our experimental manipulation supports the hypothesis that these constraints can have implications for individual sociality even on a very fine scale.

### The effect of social group stability treatment on flock size and social network centrality (H3)

We expected individuals in the stable treatment to have stronger ties with fewer individuals at the end of the assortative treatment because they had more opportunities to interact repeatedly with the same individuals compared to individuals who shared feeders with different individuals across each of the three phases. Flock size increased in both the stable and unstable social group treatments, which is to be expected since restricting access to a single feeder led to queuing, or individuals being around the feeders for longer as they determined which feeder they could access. However, against our prediction, the increase in flock size was larger in the stable treatment. We suggest that one possible mechanism for this finding is that there may have been greater synchrony in arrival times among birds in flocks in the stable treatment because, for example, social information should be more reliable, i.e., individuals in stable flocks knew which flock mates to rely on for information in the context of shared vigilance or finding the correct feeder. Other systems have shown how individuals differ in their reliability with respect to sharing information about predators and how this is linked to their network positions (Croft et al., 2009) and how learning about a food resource shapes the social network, where individuals who reliably have information become more central (Kulahci et al., 2018).

We had also predicted that individuals in the stable social group would have higher strength of connections (weighted degree and weighted eigenvector centrality) because they had more opportunity to create bonds with the same individuals, and the information about feeder choice from those individuals would be more reliable, compared to those who shared feeders with different individuals across each of the three phases. Although the presence of more individuals in the stable treatment (larger increase in flock size) may have been helpful to find the appropriate feeder (e.g. through stimulus enhancement (Heyes, 1994)), the strength of connections (weighted degree) changed in a similar way in the stable and unstable treatment. Against our prediction, there was weak evidence that the overall connectedness (weighted eigenvector centrality) decreased more for the stable than the unstable group, but in support of our hypothesis, the relationship between the dyadic associations in the two phases during the manipulation was stronger in the stable group than the unstable group. This is suggestive of a trade-off between forming stable social relationships and having larger social groups (Heathcote et al., 2017). We predicted that if the environment was too unpredictable, and led to too many changes in who individuals are feeding with (i.e., in the unstable social group treatment), this could have increased the costs of retaining previous associations. Yet, we found that despite these regular changes at the feeder level, individuals were still able to maintain their weighted degree in a similar way to birds from the stable treatment, and they maintained their previous associations as measured by dyadic interaction strength, demonstrating that fine-scale changes in resource access do not always affect social ties at a larger scale of the feeding patch. Previous studies found that, in contrast, fine-scale social disturbances weakened associations between individuals (Formica et al., 2017; Maldonado-Chaparro et al., 2018). However, those studies involved disturbances that included visually separating individuals for several days. In our study, there was no manipulation of which individuals could be a member of the social group, but we did manipulate the fine-scale access to food within the larger feeder array. This difference in methodologies may explain the difference between our results and those of previous studies: physically separating individuals for several days had stronger negative effects on social bonds between individuals than did our manipulation that merely forced individuals to forage at different micro-sites but otherwise allowed them to remain in the same social group.

### Persistence of effects (H4)

We also aimed to understand how perturbations carried over into time periods following the perturbations. Some effects of our food distribution and sociality manipulations persisted over time, even across subsequent changes in food distribution. We found that at the end of the experiment – during the final dispersed phase – individuals came to the feeders in larger flocks, had stronger associations with other individuals, and had more central associates, compared to the same dispersed configuration at the beginning of the experiment. This persistence observed in our experiment may have both spatial and temporal explanations. First, the clustered treatment had a high density of feeders, forcing individuals to interact at close range. This may have increased opportunities for social bond formation, and indeed we observed an increase in social connectedness. Once these social associations had been formed, they may have carried over into new contexts. For instance, Firth and Sheldon (2015) found that controlling access to feeders changed the social network in a foraging context not only at those feeders, but also at unrestricted feeders, and even while prospecting for nests in the context of breeding. Second, the clustered phase of the experiment was relatively long duration compared to the other phases. This would have again given individuals increased opportunities to interact and form stable relationships. Once solidified, these were then likely to continue for some time after the distribution of resources changed. However, Heinen et al. (2022), who used a similar timeline as our experiment, found that no significant assortment persisted beyond the initial manipulation. Time spent together does not necessarily influence the strength of the relationship (Boucherie et al., 2017, 2018; Proops et al., 2021), as relationships are dynamic and change overtime. However, a threshold of time spent together may be necessary to create bonds, which then take a certain time to change. This raises important questions for the study of social networks: over what time scale are social bonds formed, and how does this interact with the duration of continued social ties following an environmental disturbance?

Given the temporal set up of our experiment, it is difficult to determine which specific treatment led to those persistent effects. Flock size increased through both the physical and social manipulations throughout the experiment, suggesting that the higher flock size observed at the end is due to either additive or non-additive effects that carried over from the different manipulations to affect sociality in the final dispersed phase. Weighted degree and weighted eigenvector centrality showed increases or decreases throughout our experiment, depending on the treatment applied. It is therefore difficult to disentangle the effect of each manipulation. However, effect sizes were larger for the dispersed to clustered manipulation (H1), compared to the manipulation restricting access to feeders (H2), and the manipulation of social stability (H3). The direction and size of the effect during H1 are similar to those comparing the final to the initial dispersed phase (H4), suggesting that this initial manipulation of food distribution may have had the strongest effect of any of our manipulations in the long term. Along the same line, by the end of the experiment, we did not find any significant effects of social stability treatment on our social network metrics, except for group size, but such differences could be explained entirely by the fact that groups that were later assigned to the unstable social treatment had smaller flock sizes at the start of the experiment. This suggests that our social manipulation mostly influenced changes in social network while the manipulation occurred, but had little long-lasting effects. Likewise, Heinen et al. (2022) - unlike Firth and Sheldon (2015) - found that after assorting individuals at food patches, in a similar manner to our social stability treatment, the assortment did not persist into a new feeding context.

### Dyadic associations

We also show that dyadic associations between individuals were maintained over time. Despite the fine scale of the manipulation, and its persistent impact on social network metrics, we found evidence of consistent social ties across periods. Great tits appear to be highly consistent in their social associations: spring breeding territorial distributions reflected winter foraging network positions (Firth & Sheldon 2016), and individuals temporarily removed from the flock resumed their prior social associations upon reintroduction (Firth et al. 2017). However, such consistency is not found in all species. For instance, work in gannets (*Morus serrator*) found that the identity of associates was not consistent across different foraging contexts (Jones et al., 2020), and an experimental manipulation showed that chickadees (*Poecile gambeli*) restructure their network by assorting mostly with birds assigned to the same resource (Heinen et al., 2022). To determine the true cause of variation across species in social network stability in the face of disturbance, additional experiments in other species like the ones presented here (manipulating habitat-based and social factors) would be very valuable.

Here we show that despite both physical and social experimental changes in their environment, the social bonds individual great tits formed with conspecifics were preserved. Living in fission-fusion flocks might have selected for strategies that allow some buffering against perturbation and allow for individuals to consistently associate with the same conspecifics. This raises the question of how individuals buffer such environmental changes in their network, and whether this has consequences for their fitness and life history strategies. For instance, while associations are maintained even when individuals are assorted at different feeders, scrounging might increase as individuals still use feeders where they don’t have access to food in order to maintain previously established relationships (Regan et al., 2022). Data outside of the feeding context might also prove useful in understanding how individuals adjust their behaviour to stay with their associates despite changes in their foraging environment. Further work to better understand the underlying mechanisms from which such social stability emerges and is maintained will be important in understanding the evolutionary forces acting on social structure (Farine & Sheldon, 2019).

## Conclusion

We show that even in the face of direct fine-scale manipulations of the physical and social environment, individual differences in social behaviour (flock size and social network centralities) are maintained in a wild population of birds. We also found that dyadic social associations remained consistent under these perturbations, as individuals consistently associated with the same individuals. Such consistency is in line with previous work, and suggests personality differences in social behaviour during foraging, and some resilience of social associations to environmental change. But the environment also drives plasticity in social network metrics. We found that manipulating a combination of habitat-based and social factors can have persistent effects (i.e. beyond the initial manipulation) on social network structure. Even fine scale-changes in food distribution and interactions at a feeder can have effects on network metrics, showing that social traits are dynamic.

Our results provide a novel insight into how fine scale manipulations of socio-environmental factors have persistent effects on group structure and stability, and that relative social differences among individuals may be robust to these perturbations.

## Acknowledgements

We thank Keith McMahon for assisting with fieldwork and bird ringing; Karen Cogan for assisting with equipment sourcing and purchasing; Martin Whitaker for helping design the selective feeders; Ella Cole for logistical advice at Whytham; Ipek Kulahci, James Savage and Iván de la Hera for helping with their constructions. Ben Sheldon provided access to the main study system at Wytham Woods and funding (ERC AdG 250164) to maintain the long-term study.

