## Supplementary Tables 1-12, Supplementary Figures 1-3 for "Unravelling effects of fine-scale changes within wild-bird flocks on sociality"

I. **Methods**

Table S1: Summary data of unique individuals at each step of the experiment, including the number of males, females, adults and juveniles

| Resource treatment | Total | Adult Female | Adult Male | Adult Unknown Sex | Juvenile Female | Juvenile Male | Juvenile Unknown Sex |
| --- | --- | --- | --- | --- | --- | --- | --- |
| Initial dispersed | 198 | 60 | 71 | 4 | 23 | 30 | 10 |
| Open clustered | 167 | 52 | 58 | 3 | 18 | 25 | 11 |
| Assortative clustered |  |  |  |  |  |  |  |
| Phase I | 117 | 33 | 40 | 1 | 18 | 18 | 7 |
| Phase II | 116 | 33 | 39 | 0 | 18 | 21 | 5 |
| Phase III | 114 | 34 | 39 | 0 | 18 | 19 | 4 |
| Final dispersed | 148 | 43 | 53 | 1 | 21 | 25 | 5 |
| Total | 258 | 77 | 85 | 5 | 33 | 43 | 15 |
| Birds that took part in the initial dispersed and open clustered treatment | 131 | 42 | 48 | 2 | 13 | 18 | 8 |
| Birds that took part in the open clustered and assortative clustered P1 treatment | 107 | 29 | 40 | 1 | 15 | 16 | 6 |
| Birds that took part in all 3 assortative clustered treatments | 94 | 29 | 32 | 0 | 16 | 14 | 3 |
| Birds that took part in all 6 treatments or phases | 68 | 19 | 28 | 0 | 8 | 12 | 1 |

II. **Results: effects of food distribution and social treatment on activity, flock size and social network metrics**

Table S2: Linear mixed models showing the effect of treatment on each of four dependent variables linked to bird usage of feeders during stages or phases over the course of the experiment: the number of individuals present, the number of visits per day, and number of flocks individuals came to the feeders with. For the site level analysis (number of individuals per stage), we included site as a random effect (n=8 sites). For the individual level analysis (number of visits per individual and number of flocks individuals came in), we included site and individual identity as random effects (n=238 unique individuals). ^1^ baseline = initial dispersed; ^2^ baseline = female; ^3^ baseline = adult

| Dependent Variable | Independent variables | Estimate (SE) | 95% CI | P value |
| --- | --- | --- | --- | --- |
| Number of individuals | Intercept | 3.23 (0.143) | 2.91; 3.53 | <0.001 |
|  | Resource (open clustered)^1^ | -0.214 (0.102) | -0.416; -0.013 | 0.037 |
|  | Resource (assortative clustered P1)^1^ | 0.028 (0.096) | -0.161; 0.217 | 0.772 |
|  | Resource (assortative clustered P2)^1^ | -0.009 (0.097) | -0.200; 0.181 | 0.923 |
|  | Resource (assortative clustered P3)^1^ | -0.088 (0.099) | -0.283; 0.106 | 0.372 |
|  | Resource (final dispersed)^1^ | -0.280 (0.104) | -0.486; -0.076 | 0.007 |
| Visit Count / Number of Days in Treatment | Intercept | 43.9 (6.89) | 30.1; 57.6 | <0.001 |
|  | Sex (male)^2^ | 2.56 (4.37) | -5.96; 11.2 | 0.558 |
|  | Age (juveniles)^3^ | 0.435 (4.82) | -9.03; 9.84 | 0.928 |
|  | Resource (open clustered)^1^ | -6.37 (4.48) | -15.1; 2.49 | 0.156 |
|  | Resource (assortative clustered P1)^1^ | 29.8 (4.98) | 20.0; 40.0 | <0.001 |
|  | Resource (assortative clustered P2)^1^ | 49.8 (5.00) | 30.9; 51.0 | <0.001 |
|  | Resource (assortative clustered P3)^1^ | 55.1 (5.01) | 45.2; 65.2 | <0.001 |
|  | Resource (final dispersed)^1^ | 34.9 (4.61) | 25.9; 44.0 | <0.001 |
| Number of flocks individuals came to the feeders/ Number of days in Treatment | Intercept | 8.39 (1.51) | 5.33; 11.4 | <0.001 |
|  | Sex (male)^2^ | 0.045 (0.730) | -1.38; 1.49 | 0.951 |
|  | Age (juveniles)^3^ | -0.805 (0.807) | -2.39; 0.775 | 0.320 |
|  | Resource (open clustered)^1^ | -1.33 (0.714) | -2.73; 0.079 | 0.062 |
|  | Resource (assortative clustered P1)^1^ | 3.34 (0.792) | 1.77; 4.96 | <0.001 |
|  | Resource (assortative clustered P2)^1^ | 3.58 (0.797) | 2.00; 5.20 | <0.001 |
|  | Resource (assortative clustered P3)^1^ | 6.50 (0.798) | 4.92; 8.12 | <0.001 |
|  | Resource (final dispersed)^1^ | 7.99 (0.735) | 6.56; 9.44 | <0.001 |

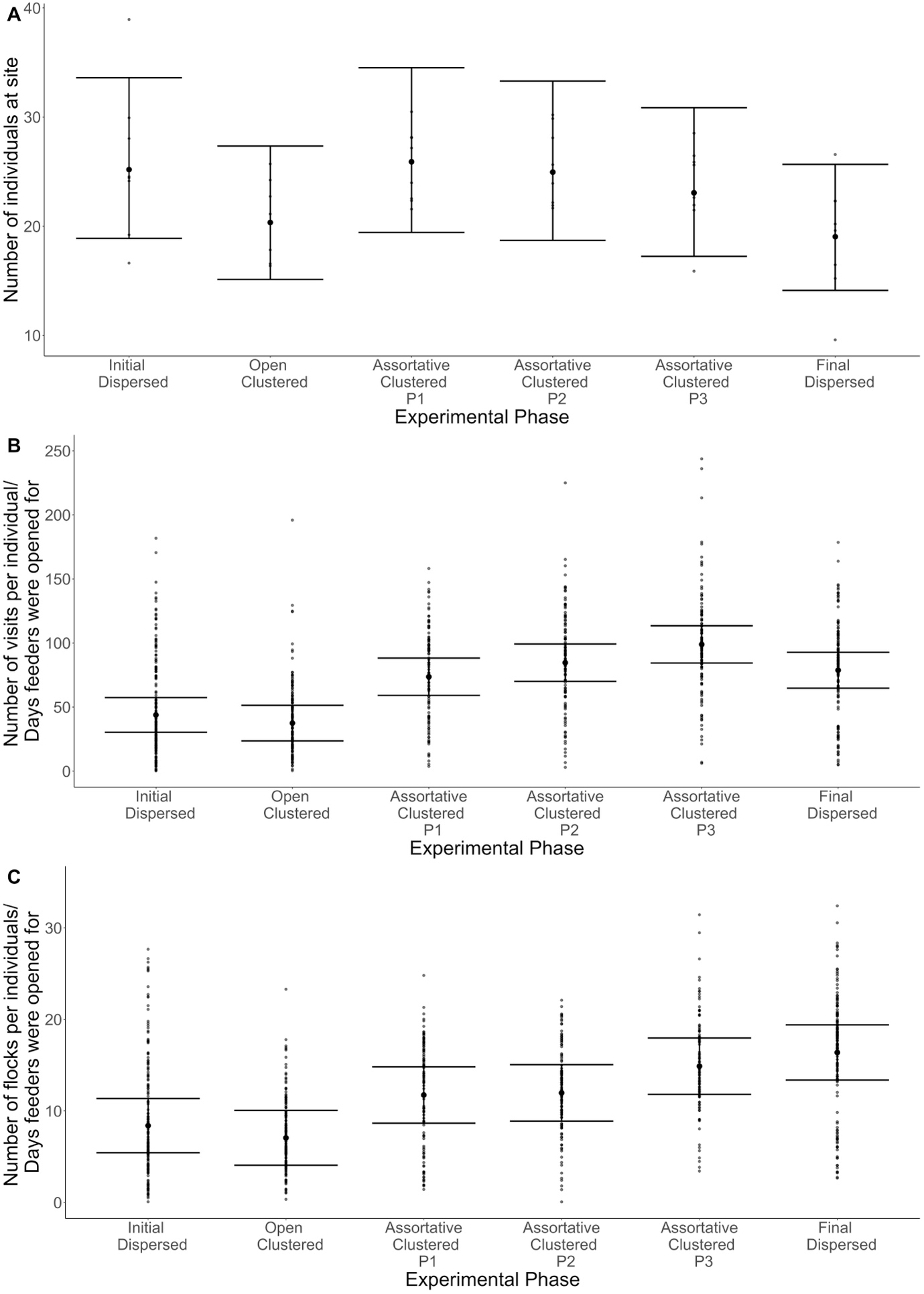
Figure S1: Partial residual plots showing the relationship between experimental stage and (A) number of individuals at sites, (B) number of visits per individual per day, and (C) number of flocking events per individual per day, across the initial dispersed, open clustered, assortative clustered P3, and final dispersed treatment levels. Error bars are 95% confidence intervals

**H1: How does changing food distribution from dispersed to clustered influence social behaviour?**

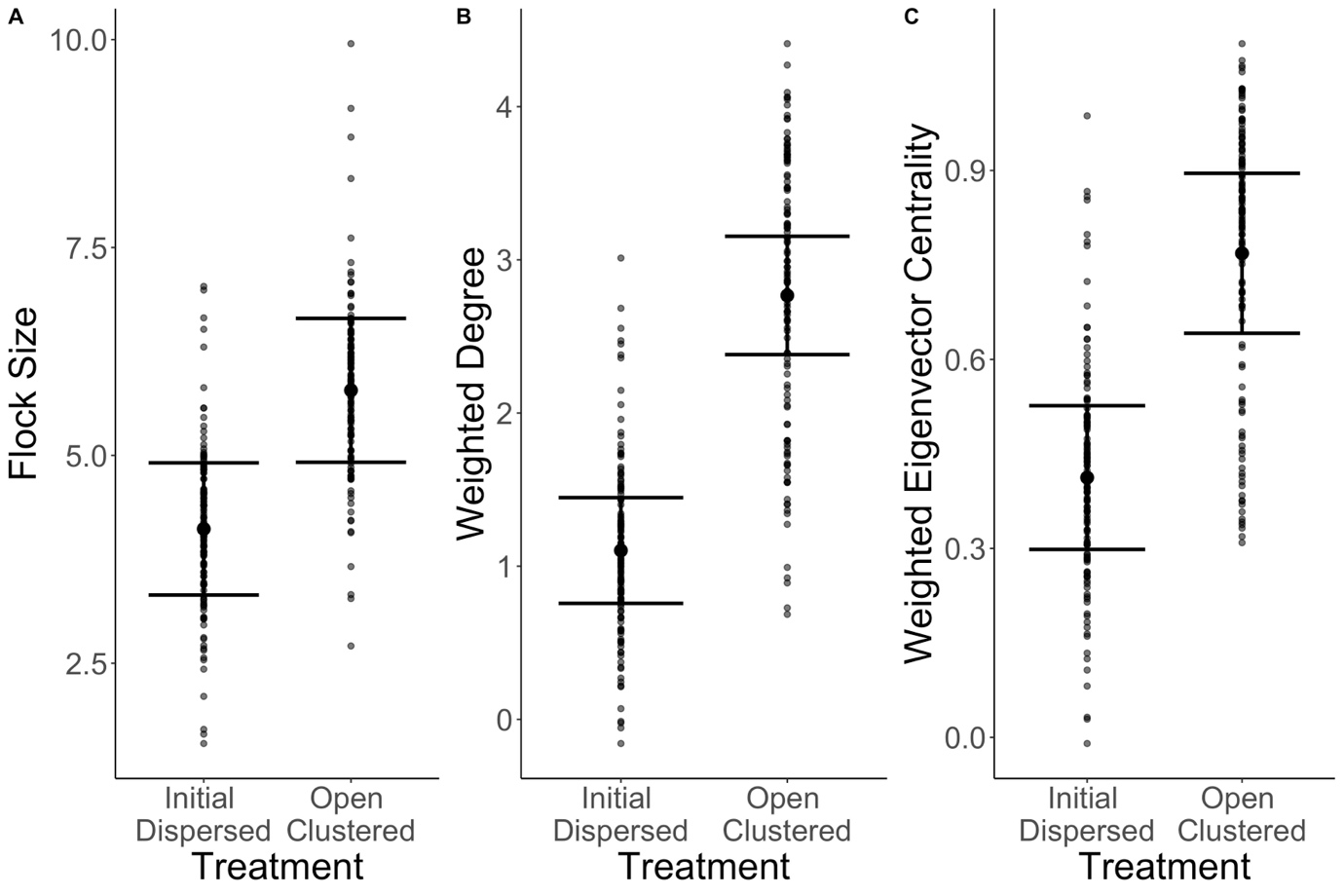
Figure S2: Partial residual plots showing changes in (A) flock size, (B) weighted degree and (C) weighted eigenvector centrality when food first changed from dispersed to clustered treatments. Error bars are 95% confidence intervals based on model from Table 1.

Table S3: Linear mixed models of how each of four individual social network metrics changed from the initial dispersed to the open clustered treatments. Site and individual identity were included as random effects. ^1^ baseline = female; ^2^ baseline = adult; ^3^ baseline = initial dispersed

| Dependent variable | Independent variables | Estimate (SE) | 95% CI | P value |
| --- | --- | --- | --- | --- |
| Flock Size | Intercept | 2.24 (0.831) | 0.609; 3.90 | 0.012 |
|  | Flocking events | -0.002 (0.002) | -0.005; 0.001 | 0.257 |
|  | Individuals in local network | 0.072 (0.026) | 0.020; 0.123 | 0.008 |
|  | Sex (male)^1^ | -0.092 (0.136) | -0.356; 0.173 | 0.499 |
|  | Age (juveniles)^2^ | -0.046 (0.158) | -0.352; 0.263 | 0.772 |
|  | Resource (open clustered)^3^ | 1.67 (0.224) | 1.23; 2.12 | <0.001 |
| Weighted Degree | Intercept | -1.020(0.449) | -2.14; -0.313 | 0.015 |
|  | Flocking events | 0.013 (0.001) | 0.011; 0.016 | <0.001 |
|  | Individuals in local network | 0.054 (0.014) | 0.025; 0.085 | 0.002 |
|  | Sex (male)^1^ | -0.003 (0.099) | -0.196; 0.191 | 0.977 |
|  | Age (juveniles)^2^ | 0.014 (0.115) | -0.209; 0.239 | 0.905 |
|  | Resource (open clustered)^3^ | 1.66 (0.143) | 1.38; 1.94 | <0.001 |
| WEVC | Intercept | -0.162 (0.137) | -0.445; 0.118 | 0.252 |
|  | Flocking events | 0.005 (0.0003) | 0.004; 0.006 | <0.001 |
|  | Individuals in local network | 0.010 (0.004) | 0.001; 0.019 | 0.037 |
|  | Sex (male)^1^ | -0.0005 (0.026) | -0.053; 0.050 | 0.985 |
|  | Age (juveniles)^2^ | -0.034 (0.031) | -0.062; 0.058 | 0.912 |
|  | Resource (open clustered)^3^ | 0.356 (0.041) | 0.271; 0.435 | <0.001 |

Table S4: Permutation analyses of models in Table S3, which examined the changes in each of the four individual social network metrics, from the initial dispersed to the open clustered resource treatment. p values show where the observed estimates fall within the distribution of estimates from the permutations. ^1^ baseline = female; ^2^ baseline = adult

| Dependent variable | Intercept | Nb flocking events | Nb individuals in local network | Sex (male)^1^ | Age (juvenile)^2^ | Initial dispersed vs open clustered resources |
| --- | --- | --- | --- | --- | --- | --- |
| Flock size | 0.000 | 0.242 | 0.004 | 0.494 | 0.774 | 0.000 |
| Weighted degree | 0.000 | 0.000 | 0.000 | 0.982 | 0.878 | 0.000 |
| WEVC | 0.000 | 0.000 | 0.016 | 0.946 | 0.960 | 0.000 |

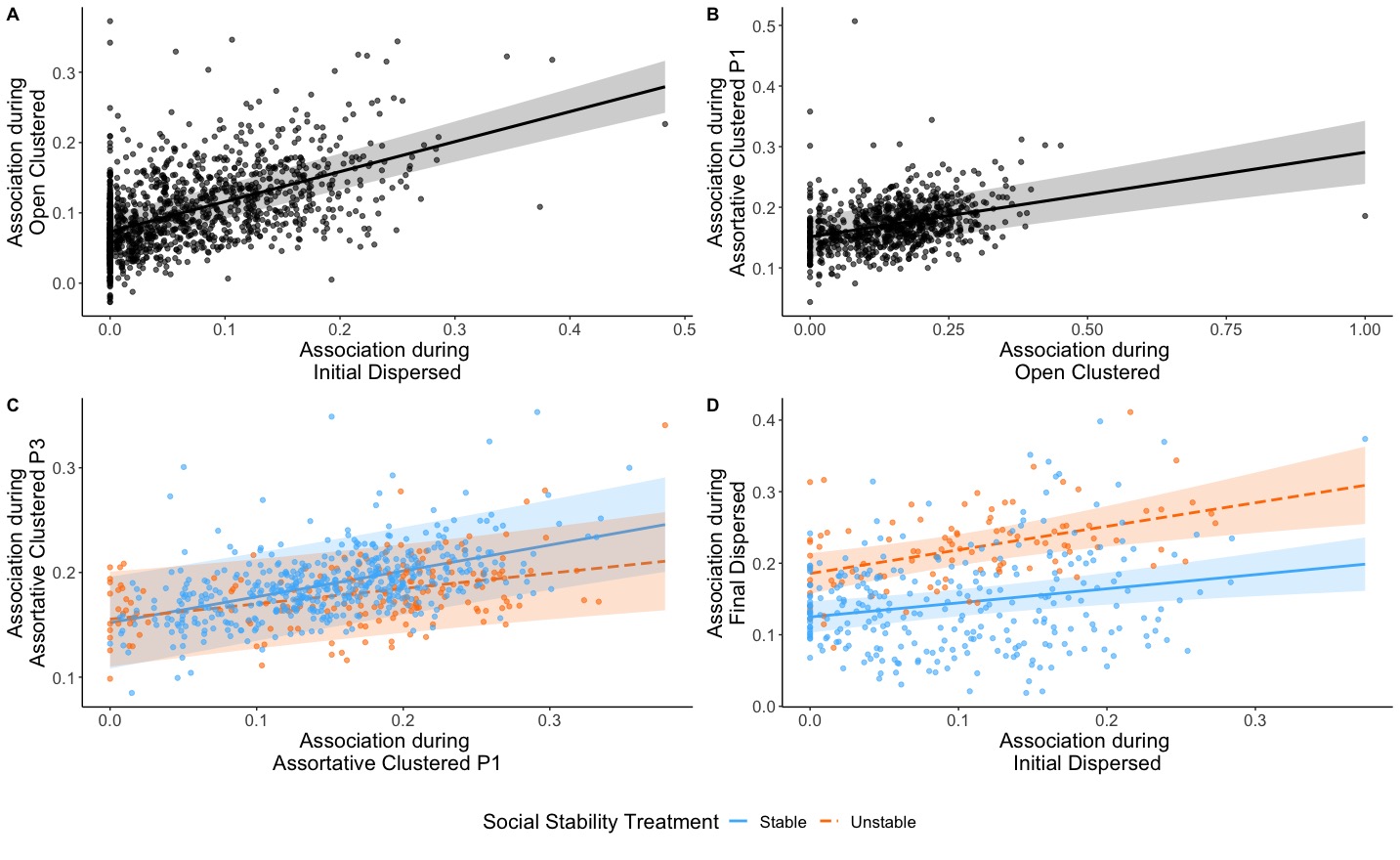
Figure S3: Partial residual plots showing how pairwise associations in one treatment predicted these in the next for: a) the initial dispersed treatment and the open clustered treatment; b) the open clustered treatment and the assortative clustered treatment (P1); c) the assortative clustered P1 treatment and P3 treatment; and d) the initial dispersed and final dispersed treatments. For c) and d), separate lines are shown for birds in the stable (blue) and unstable (orange) social stability treatments. Interaction was ns. for c) and d). Shaded areas are the 95% confidence intervals from corresponding models in the main text and in Table S7 and S10. We added random effects for the identity of both individuals of each dyad and site.

**H2: Does manipulated assortative-feeding on a fine spatial scale change social behaviour?**

Table S5: Permutation analyses of models in Table 1, which examined the changes in each of the four individual social network metrics, from the open clustered to the assortative clustered P1 resource treatment. p values show where the observed estimates fall within the distribution of estimates from the permutations. ^1^ baseline = female; ^2^ baseline = adult

| Dependent variable | Intercept | Nb flocking events | Nb individuals in local network | Sex (male)^1^ | Age (juvenile)^2^ | Open clustered vs assortative clustered |
| --- | --- | --- | --- | --- | --- | --- |
| Flock size | 0.000 | 0.594 | 0.000 | 0.400 | 0.818 | 0.000 |
| Weighted degree | 0.000 | 0.000 | 0.000 | 0.866 | 0.390 | 0.000 |
| WEVC | 0.584 | 0.000 | 0.164 | 0.944 | 0.044 | 0.000 |

**H3: Does social stability influence the effect of the assortative-feeding manipulation on social behaviour?**

Table S6: posthoc analysis of the interaction between resource and social stability treatment from the model in Table 2, showing the pairwise difference between the assortative clustered P1 and P3 phases

| Dependent variable | Social treatment | Estimate | SE | 95% C.I. | p-value |
| --- | --- | --- | --- | --- | --- |
| Flock size | Stable | -1.02 | 0.097 | -1.21; -0.830 | <0.001 |
|  | Unstable | -0.424 | 0.114 | -0.647; -0.201 | <0.001 |
| Weighted degree | Stable | -0.232 | 0.057 | -0.343; -0;121 | <0.001 |
|  | Unstable | -0.225 | 0.066 | -0.355; -0.095 | 0.001 |
| WEVC | Stable | 0.099 | 0.020 | 0.061; 0.138 | <0.001 |
|  | Unstable | 0.050 | 0.023 | 0.005; 0.095 | 0.031 |

Table S7: Permutation analyses of models in Table 2, which examined how changes in each of the four individual social network metrics, from the start (P1) to the end (P3) of the assortative clustered resource treatment level, were influenced by social stability, as tested by their interaction. p values show where the observed estimates fall within the distribution of estimates from the permutations. ^1^ baseline = female; ^2^ baseline = adult

| Dependent variable | Intercept | Nb flocking events | Nb individuals in local network | Sex (male)^1^ | Age (juvenile)^2^ | Clustered P1 vs clustered P3 | Social stability treatment | Interaction |
| --- | --- | --- | --- | --- | --- | --- | --- | --- |
| Flock size | 0.000 | 0.522 | 0.000 | 0.836 | 0.634 | 0.000 | 0.648 | 0.000 |
| Weighted degree | 0.000 | 0.000 | 0.002 | 0.348 | 0.356 | 0.000 | 0.282 | 0.948 |
| WEVC | 0.092 | 0.000 | 0.000 | 0.654 | 0.642 | 0.000 | 0.186 | 0.118 |

Table S8: Model examining the relationship between dyadic associations during the third phase (P3) and the first phase (P1) of the assortative clustered treatment level, and whether this varied depending on social stability treatment. Site and individual identity of both individuals of each dyad were included as random effects.^1^ baseline = stable

| Factor | Estimate (SE) | 95% CI | P value |
| --- | --- | --- | --- |
| Intercept | 0.152 (0.022) | 0.109; 0.194 | <0.001 |
| Dyadic association during P1 | 0.248 (0.042) | 0.166; 0.330 | <0.001 |
| Social stability (unstable)^1^ | 0.003 (0.032) | -0.057; 0.065 | 0.919 |
| Dyadic association during P1 × social stability (unstable)^1^ | -0.101 (0.065) | -0.228; 0.031 | 0.121 |

**H4: Were the changes observed during the clustered treatment persistent when dispersed food treatment was restored?**

Table S9: Linear mixed models that examined how changes in each of the four individual social network metrics, from the initial dispersed to the final dispersed resource treatment, were influenced by social stability, as tested by their interaction. Site and individual identity were included as random effects. ^1^ baseline = female; ^2^ baseline = adult; ^3^ baseline = initial dispersed; ^4^baseline = stable

| Dependent variable | Independent variables | Estimate (SE) | 95% CI | P value |
| --- | --- | --- | --- | --- |
| Flock Size | Intercept | 1.48 (0.695) | 0.187; 2.77 | 0.050 |
|  | Flocking events | 0.00002 (0.001) | -0.003; 0.003 | 0.990 |
|  | Individuals in local network | 0.107 (0.017) | 0.074; 0.139 | <0.001 |
|  | Sex (male)^1^ | -0.028 (0.128) | -0.274; 0.222 | 0.831 |
|  | Age (juveniles)^2^ | 0.121 (0.140) | -0.146; 0.394 | 0.391 |
|  | Resource (final dispersed)^3^ | 1.85 (0.162) | 1.54; 2.16 | <0.001 |
|  | Social stability (unstable)^4^ | -1.08 (0.774) | -2.54; 0.395 | 0.209 |
|  | Resource (final dispersed)^3^ × social stability (unstable)^4^ | 1.26 (0.344) | 0.606; 1.93 | <0.001 |
| Weighted Degree | Intercept | -0.852 (0.353) | -1.50; -0.192 | 0.023 |
|  | Flocking events | 0.010 (0.001) | 0.009; 0.012 | <0.001 |
|  | Individuals in local network | 0.051 (0.010) | 0.031; 0.070 | <0.001 |
|  | Sex (male)^1^ | -0.009 (0.080) | -0.163; 0.146 | 0.908 |
|  | Age (juveniles)^2^ | 0.015 (0.087) | -0.151; 0.185 | 0.861 |
|  | Resource (final dispersed)^3^ | 1.71 (0.101) | 1.52; 1.91 | <0.001 |
|  | Social stability treatment (unstable)^4^ | -0.156 (0.336) | -0.788; 0.471 | 0.657 |
|  | Resource (final dispersed)^3^ × social stability treatment (unstable)^4^ | 0.281 (0.210) | -0.139; 0.672 | 0.184 |
| WEVC | Intercept | 0.241 (0.096) | 0.062; 0.418 | 0.020 |
|  | Flocking events | 0.004 (0.0002) | 0.004; 0.005 | <0.001 |
|  | Individuals in local network | -0.004 (0.003) | -0.009; 0.002 | 0.190 |
|  | Sex (male)^1^ | -0.007 (0.022) | -0.050; 0.035 | 0.757 |
|  | Age (juveniles)^2^ | 0.016 (0.024) | -0.031; 0.062 | 0.500 |
|  | Resource (final dispersed)^3^ | 0.278 (0.028) | 0.224; 0.332 | <0.001 |
|  | Social stability treatment (unstable)^4^ | 0.089 (0.091) | -0.084; 0.257 | 0.365 |
|  | Resource (final dispersed)^3^ × social stability treatment (unstable)^4^ | -0.072 (0.058) | -0.180; 0.041 | 0.213 |

Table S10: Permutation analyses of models in Table S8, which examined how changes in each of the four individual social network metrics, from the initial dispersed to the final dispersed treatments, were influenced by social stability, as tested by their interaction. P values show where the observed estimates fall within the distribution of estimates from the permutations. ^1^baseline = female; ^2^baseline = adult; ^3^baseline = initial dispersed; ^4^baseline = stable.

| Dependent variable | Intercept | Nb flocking events | Nb individuals in local network | Sex (male)^1^ | Age (juvenile)^2^ | Resource (final dispersed)^3^ | Social stability (unstable)^4^ | Resource (final dispersed)^3^ × Social stability (unstable)^4^ |
| --- | --- | --- | --- | --- | --- | --- | --- | --- |
| Flock size | 0.000 | 0.978 | 0.000 | 0.824 | 0.392 | 0.000 | 0.144 | 0.000 |
| Weighted degree | 0.000 | 0.000 | 0.000 | 0.886 | 0.854 | 0.000 | 0.690 | 0.230 |
| WEVC | 0.002 | 0.000 | 0.178 | 0.728 | 0.490 | 0.000 | 0.296 | 0.204 |

Table S11: posthoc analysis of the interaction between resource and social stability treatment from the model in Table S8, showing the pairwise difference between the initial and final dispersed treatment

| Dependent variable | Social treatment | Estimate | SE | 95% C.I. | p-value |
| --- | --- | --- | --- | --- | --- |
| Flock size | Stable | -1.85 | 0.163 | -2.16; -1.53 | <0.001 |
|  | Unstable | -3.11 | 0.317 | -3.73; -2.49 | <0.001 |
| Weighted degree | Stable | -1.71 | 0.102 | -1.91; -1.51 | <0.001 |
|  | Unstable | -1.99 | 0.193 | -2.36; -1.61 | <0.001 |
| WEVC | Stable | -0.278 | 0.028 | -0.333; -0.223 | <0.001 |
|  | Unstable | -0.206 | 0.053 | -0.310; -0.102 | <0.001 |

Table S12: Mixed model examining the relationship between dyadic associations during the final dispersed phase and the initial dispersed phase of the resource treatment, and whether this varied depending on social stability treatment. Site and individual identity of both individuals of each dyad were included as random effects. ^1^ baseline = stable

| Factor | Estimate (SE) | 95% CI | P value |
| --- | --- | --- | --- |
| Intercept | 0.125 (0.011) | 0.104; 0.146 | <0.001 |
| Dyadic associations during initial dispersed | 0.197 (0.059) | 0.081; 0.321 | 0.001 |
| Social stability (unstable)^1^ | 0.061 (0.018) | 0.027; 0.094 | 0.007 |
| Dyadic associations during initial dispersed × Social stability (unstable) ^1^ | 0.132 (0.109) | -0.078; 0.349 | 0.229 |
